# Optimal virulence in ageing populations

**DOI:** 10.64898/2026.03.19.712865

**Authors:** Jessica Clark, Luke McNally, Tom Little

**Affiliations:** School of Biodiversity, One Health & Veterinary Medicine, University of Glasgow, Scotland, UK; Institute of Ecology & Evolutionary Biology, University of Edinburgh, Scotland, UK; Centre for Engineering Biology, University of Edinburgh, Scotland, UK

**Keywords:** Virulence evolution, healthspan, demography, infectious diseases, trade-offs

## Abstract

Global populations are ageing at an unprecedented rate. For many diseases, age is a strong indicator of susceptibility, morbidity, or mortality. Principles of evolutionary biology can be leveraged to understand how pathogens may optimally exploit new populations, and the impact of this on the global burden of infectious-disease-induced mortality. We parameterised an age-specific R_0_ model with 2017 epidemiological data on Measles, Tuberculosis, Meningitis, and Ebola, and age-specific demographic estimates for 2017 and 2050, for the seven Global Burden of Disease super-regions. We explored the theoretical trade-offs between pathogen virulence & transmission, and virulence & host recovery, parameterising trade-off parameters using Latin Hypercube Sampling. Population ageing between 2017-2050 saw an increase in virulence induced mortality in four settings: 1) Ebola in sub-Saharan Africa, 2) Measles in central/eastern Europe & central Asia region, 3) Measles in North Africa & the Middle East and 4) Tuberculosis in the central/eastern Europe & central Asia region. The decrease in infection duration due to an increase of elderly people drives pathogen virulence down for diseases in the remaining settings. Understanding the mechanisms that shape pathogen dynamics and leveraging this to predict future challenges is key to endemic disease management in a rapidly changing world.

**Author Summary:** Key aspects of disease transmission including susceptibility to infection, the severity of infection, and the probability of dying from that infection, are affected by host age. Global populations are rapidly ageing, so that the mean age of most populations is generally higher than it used to be and is set to continue on this trajectory. This suggests that the dynamics of infectious diseases are also likely to change, although infectious disease dynamics tend to be non-linear as these key parameters interact. We have developed a dynamic modelling framework to explore how changes in population age structure might impact the optimal level of pathogen virulence in a population. We have chosen four infectious diseases as case studies, that differentially impact certain age classes to illustrate these dynamics. We have parameterised this framework with open access data for each of the seven Global Burden of Disease super-regions and show that population ageing can increase virulence for several diseases in differing global regions, whilst increased background rates of mortality can drive virulence down in others.

## Introduction

For the first time in human history those aged over 65 outnumber those under the age of five, and between 2015-2030 the proportion of the population aged over 65 is set to increase by 60%(1). Individuals of different ages commonly differ in their susceptibility to or ability to clear a pathogenic challenge. They also differ in other key aspects of epidemiology, like extrinsic and intrinsic rates of mortality. These differences were made abundantly clear during the Covid-19 pandemic, where the elderly were more at risk of developing severe morbidities or death. When an individual is infected, a pathogen must replicate at sufficient levels to transmit to the next. However, replication cannot be so successful that the individual is unable to recover and succumbs rapidly to disease induced mortality, which would shorten the infectious period, therefore reducing pathogen fitness. This balance between the costs and benefits of host exploitation is the basis of virulence evolution theory, which predicts a pathogen will evolve towards an optimal point of virulence, where this point may not necessarily be avirulence, but an intermediate level of host exploitation(2). This adaptive ability of pathogens is commonly observed (e.g., antimicrobial resistance) highlighting the importance of considering pathogen evolution in the face of rapid global change(3).

Most populations display some form of heterogeneity, which has been shown to influence the evolution of pathogen virulence. For example, heterogeneities in a population for factors linked to infectious disease dynamics (i.e., susceptibility or tolerance to infection), can interact with characteristics of population structure such as sex ratio, to accelerate virulence evolution (4). Optimal virulence may also be modified by population contact rates or spatial structures (5, 6) or sex ratio and sex-based heterogeneities in susceptibility or transmission routes (4, 7). From this it is evident that population structure and the differences between hosts are integral factors directing a pathogen’s optimal virulence. However, a common source of population heterogeneity that has received comparatively less attention is the effect of population age structure.

In addition to selective pressures, optimal virulence is believed to be shaped by trade-offs between transmission and pathogen virulence, and between host recovery and virulence. The transmission – virulence trade-off suggests that those pathogens producing propagules at a higher rate, transmit more rapidly, but at the expense of killing their host more quickly. Alternatively, the recovery – virulence trade off suggests that there is a cost to not being virulent enough in that a pathogen producing fewer propagules, will be cleared effectively and quickly by the host immune system, reducing the amount of time exploiting the host to maximise fitness. In terms of ageing populations, a population of older individuals will see a shorter remaining lifespan and age-specific infection induced mortality which in turn will impact the duration of infection. Coupled with the selective pressures associated with age specific susceptibility and clearance, it is likely population ageing will provide sufficient selective pressures to alter pathogen virulence.

To illustrate our hypothesis, we present a model of optimal virulence [14] under the simplest conditions, where optimal virulence is shaped only by the selective pressure of population age structure and the transmission/clearance – virulence trade-offs. Whilst there is a clear understanding that many diseases are impacted by age-structured contacts (8), it is not the ambition of this manuscript to investigate this mechanism, nor to recapitulate the full dynamics of transmission. We therefore focus on characterising the sensitivity and possible variation of pathogen virulence due to age-specific host characteristics such as susceptibility and mortality (baseline and disease induced), parameterising our model framework using publicly available demographic and infectious disease data representing the Global Burden of Disease’s seven Super Regions. We used publicly available demographic data (present and projections) to assess how estimated changes in population age structure in each region could impact the optimal intrinsic pathogen virulence in a selection of pathogens of global concern, with differing disease induced mortality rates and age-structured presentation.

## Methods

### The model

A parasites basic reproductive rate *R*_*0*_ links the numbers of susceptible and infected individuals to determine a pathogens infectivity and epidemic-potential when introduced to a totally naïve population. If > 1, then the pathogen will spread, if <1, the pathogen will die out. As per Frank (1992) [14] its standard form is

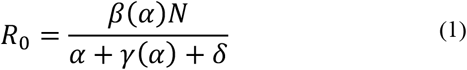

where *β*(*α*) is the transmission coefficient, *α* describes virulence and *N* the density of the population. In the denominator, *δ* is the baseline population mortality rate in the absence of the pathogen, *γ* the host recovery rate, here a function of virulence also.

Here, we adapt Frank 1992 (13) for a single infection and adapt virulence such that *α* = 𝒳*ν* where virulence is the product of host vulnerability to mortality from the pathogen, *ν*, and the parasites intrinsic virulence, 𝒳, that scales host vulnerability.

In the presence of a transmission – intrinsic virulence trade-off, as per Frank (1992) (13)

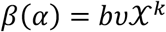

where *b* is given by *b* = *cτ* with *c* as the contact rate, and *τ* the susceptibility to infection given that contact. 𝒳 is scaled by *k*, the virulence exponent, which determines the rate of change in transmission with changing intrinsic pathogen virulence (13). *β* therefore, increases with increasing *α* until a theorised optimal point at which host exploitation by the pathogen results in sufficient damage to the host to reduce the reproductive gain of the pathogen. This is believed to lead to a trade-off between host survival and pathogen transmission.

In the presence of a trade-off between host recovery *γ* and intrinsic pathogen virulence

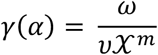

with *ω* describing the rate of pathogen clearance by the host and the virulence exponent *m* determining the optimal pathogen virulence (13). Simplified, this yields

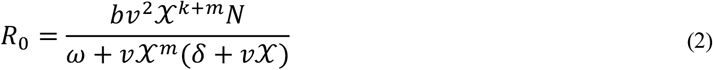

To illustrate the impact of trade-offs on optimal virulence, we assume that the only trade-off is between the transmission and virulence, such that the evolutionarily stable intrinsic virulence is *X*^∗^. The optimal infectivity at which transmission (*R*_0_) is maximised with respect to 𝒳, is then obtained by solving for 𝒳 when 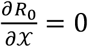, which yields

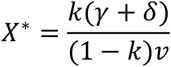

By including *ν*, but assuming a single infecting genotype, we get a similar expression to Frank (1992)(13). Any changes in virulence can therefore be determined as follows. The present evolutionary stable intrinsic virulence 𝒳 is given by

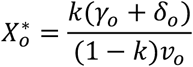

Where the subscript *o* denotes baseline values of age-specific recovery, mortality and host vulnerability respectively and

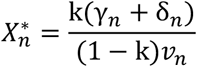

gives the new evolutionary stable intrinsic virulence, where *n* represents projected age-specific values of recovery, mortality and EV respectively, then the Δ𝒳 = 𝒳_+_ − 𝒳_∗_ so that

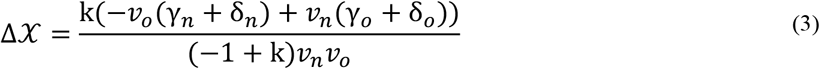

Which is positive when *k* < 1 and 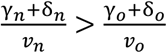

In the presence of population age structure, we assume host vulnerability, *v*, population mortality rate *δ*, recovery *γ*(*α*), and the density of susceptible hosts to be a function of host age, so that age specific pathogen 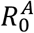 with both transmission – virulence, and recovery – virulence trade-offs is given by

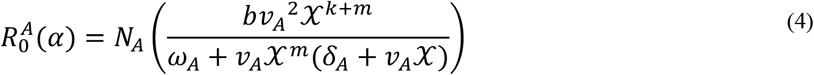

where *N*_*A*_ is the proportion of individuals in age class *A* with the total population *R*_0_ given by

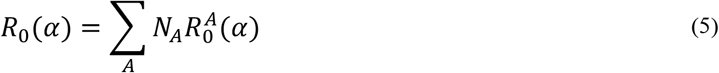

### Parameterisation

To determine how changing population age structure will influence pathogen virulence evolution, we used publicly available global infection, and demographic data to parameterise the above framework (Table 1). Countries were allocated to regions as per the Global Burden of Disease Super-regions; 1) High income, 2) Southeast Asia, East Asia, and Oceania, 3) Central Europe, Eastern Europe, and Central Asia, 4) Sub-Saharan Africa, 5) Latin America and Caribbean 6) South Asia and 7) North Africa and Middle East. Regional age specific mortality rates were provided by The World Health Organization (WHO) (Figure 1A). Age specific population density for 2017 and the 2050 projection were obtained from The United States Census Bureau and were used as the proportion of the total population made up by each 5-year age bracket (Figure 1B-H). All rates were per 100,000 people for 2017 and were normalised.

**Table 1.** Model parameter definitions, notation, and numerical data inputs.

| *Parameter* | *Notation* | *Values* |
| --- | --- | --- |
| *Population baseline mortality* | $\delta$ | *Age specific mortality* |
| *Susceptibility* | $\tau$ | *Age specific normalised incidence* |
| *Population density* | $N$ | *Age specific population density* |
| *Vulnerability* | $v$ | $\frac{\text{disease induced mortality}}{\text{incidence rate}}$ |
| *Clearance* | $\gamma$ | $\frac{1}{\frac{\text{prevalence}}{\text{incidence}}}$ |
| *Intrinsic Pathogen virulence* | $\mathcal{X}$ | *Estimated with Latin hypercube sampling.* |
| *Contact* | $c$ | *Measles = 1.2(9)*<br>*Meningitis = 0.05(10)*<br>*Tuberculosis = 0.2 (11)*<br>*Ebola = 0.15(12)* |
| *Transmission coefficient* | $b$ | *Contact x susceptibility* |
| *Virulence exponent* | $\kappa$ | *Estimated with Latin hypercube sampling* |
| *Clearance exponent* | $m$ | *Estimated with Latin hypercube sampling* |

**Figure 1.**
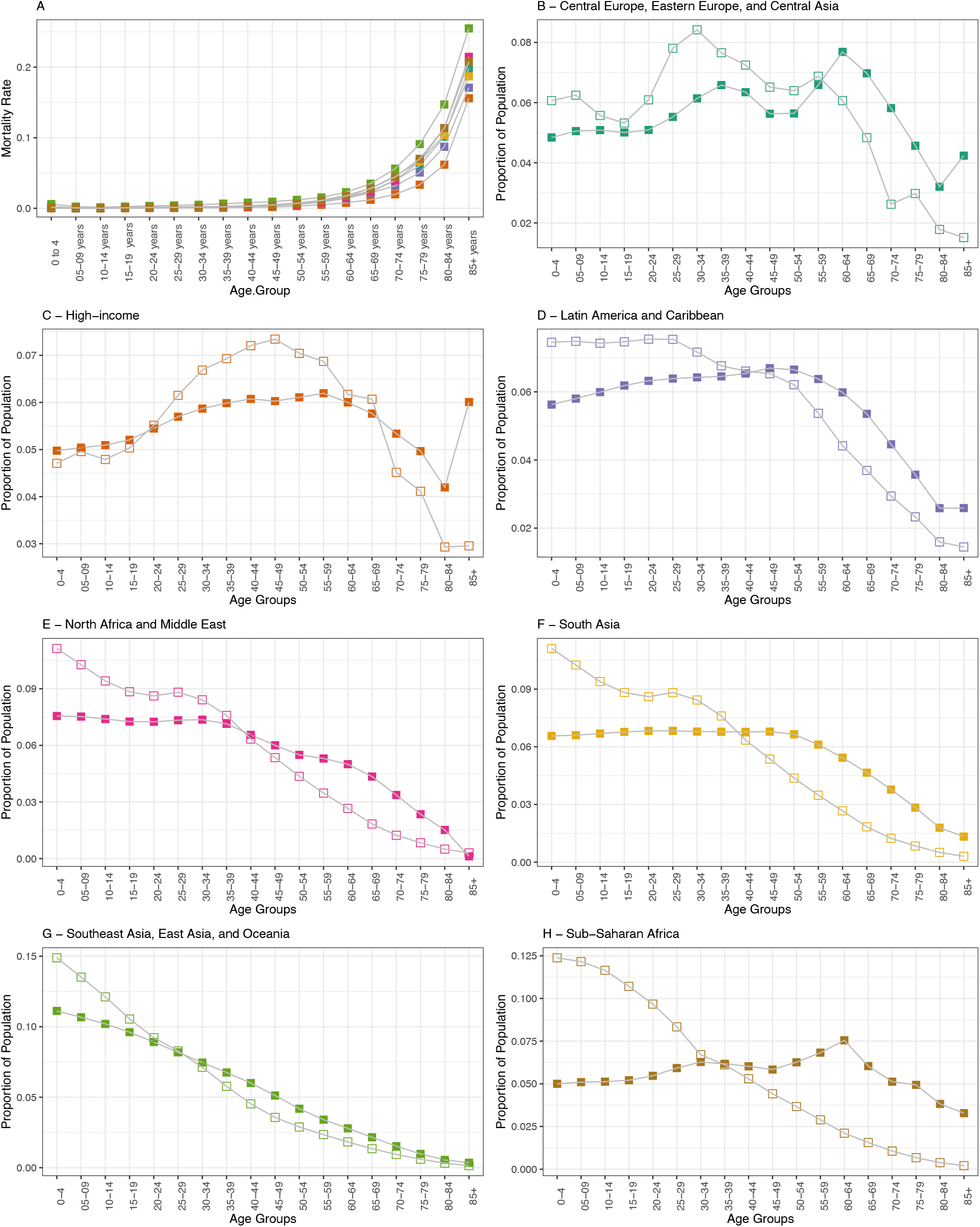
A. Age-specific rates of mortality across the seven super-regions recognised by the Global Burden of Disease. B-H. Population age structure in 2017 (open boxes) and 2050 (closed boxes) for the seven super regions recognised by the Global Burden of Disease.

To ensure all epidemiological data were from the same source and therefore processed in the same manner, we chose infectious diseases that were all present on the Global Burden of Disease database. Vector-borne diseases were not included because of the need to include both vector and host in the *R*_0_ term. We also did not include helminth infections because *R*_*0*_ relates to the number of offspring that go on to mature and reproduce from a single female (14). We therefore chose the causative agents of Ebola, bacterial Meningitis, Tuberculosis (TB), and Measles. For each disease we obtained rates for disease induced mortality (Figure S2), incidence, and prevalence per five-year age group. We note that whilst evidence in support of trade-offs is hard to acquire, previous work has investigated such trade-offs and the drivers of virulence optimisation for each pathogen (15-18).

The variables and their parameter sources can be found in Table 1. Briefly, regional age specific susceptibility to infection, *τ*, for each infection was parameterised using the normalised incidence values (Figure 2). It is evident here that whilst TB (for example) varies by location, Measles normalised incidence is strongly age-dependent but differs little across regions. Susceptibility multiplies the contact constant, *c*, to form the transmission coefficient, *b*. There were no available and unified values to describe age specific average rates of clearance for these diseases. Clearance was therefore derived from incidence and prevalence data. The average duration of infection was calculated for each region and disease as 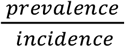 so that the rate of clearance for each region and disease could be considered 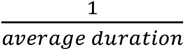. The realised virulence, *α*, in the model is denoted 𝒳*v*, capturing the relationship between the display of host response to a pathogen, *v*, and the intrinsic pathogen virulence from the parasite exploiting the host 𝒳. *v* can be considered an individual’s intrinsic age-specific risk or vulnerability to pathogen induced mortality 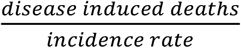

**Figure 2.**
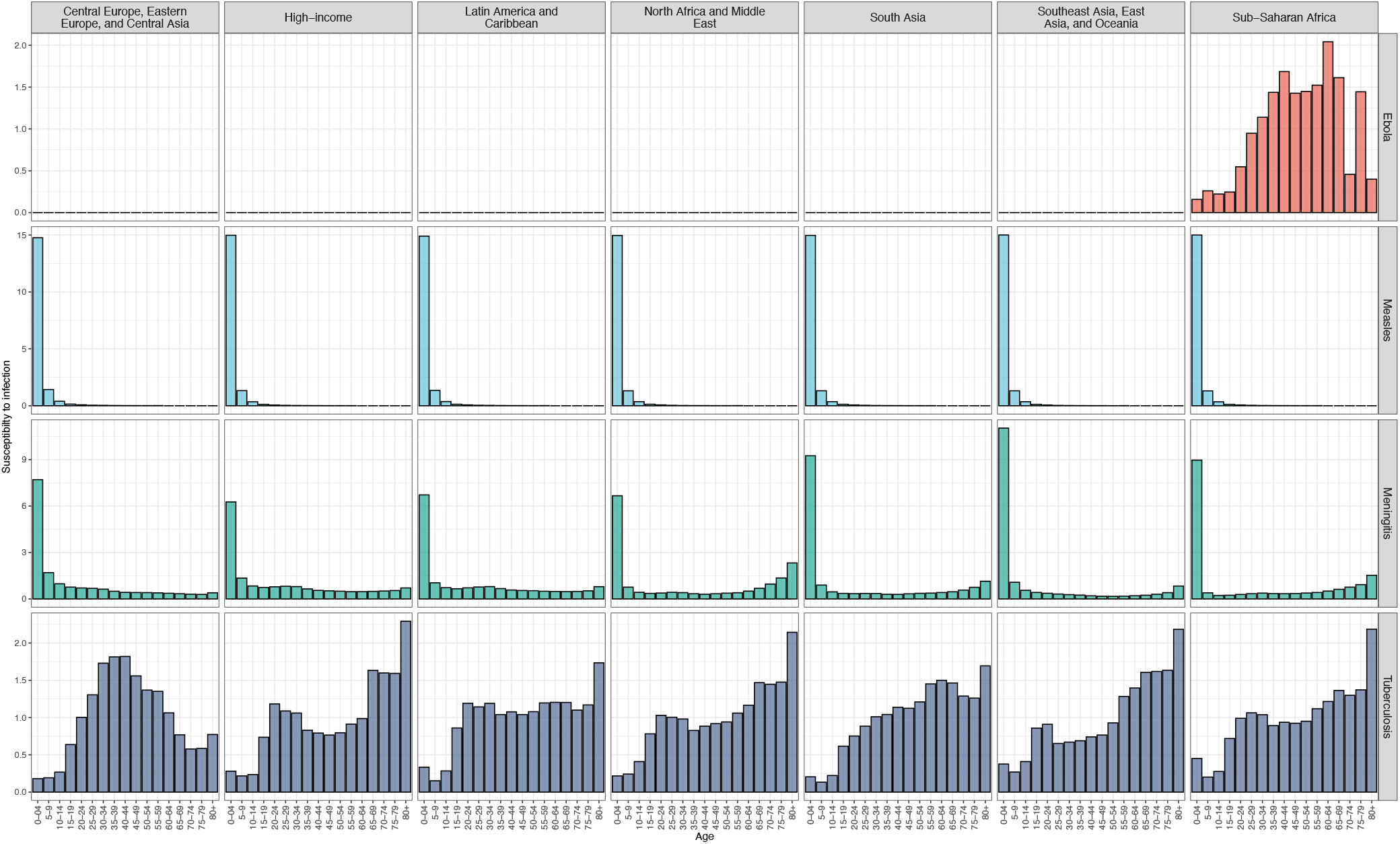
Age, region, and disease specific susceptibility to infection (normalised incidence for each age class, showing regional variation in relationship to population age structure, and variation between diseases within regions.

The final parameters of the model are the trade-off parameters: *κ* for the transmission-virulence trade off exponent, *m* for the recovery – virulence trade off, and 𝒳 for intrinsic pathogen virulence. To obtain the optimal combination of these that maximised present *R*_0_(*α*) we used Latin hyper-cube sampling(19) to explore the parameter space. As per Frank (1992) (13)*m* and *k* are bound [0,1]. As 𝒳 scales the host vulnerability to disease-induced mortality this is also bound [0,1]. We produced 1,000,000 parameter sets to explore and obtain the optimal value at which the trade-offs manifest and *R*_0_(*α*) exhibits diminishing returns. These parameter values for *k* and *m* were then used to parameterise the 2050 age structure projection, where the optimal 𝒳 was found again using the same method, but with fixed *m* and *k* from the present-day simulations (i.e., making the assumption that the only change to intrinsic virulence comes from intrinsic virulence responding to population age structure, and that the shape of the trade-offs remain the same). The LHS process for the future calculation was repeated 10,000 times to verify the solution for optimal 𝒳 and subsequent *R*_0_. Outcomes from this are shown in Table S1. Due to very little variation, we show just the mean value.

All simulations were conducted in R V4.1.3 (2022-03-10) (20) using the mclapply function from the *parallel* package (base R). All code and datafiles are available at https://github.com/iamjessclark/evolution_of_virulence.git.

## Results

### Solving for trade-off parameters

The trade-off scaling parameter *m* (Figure 3, purple)– controlling the trade-off between virulence and recovery – is more variable across diseases and regions (i.e., age structures) than the trade-off between transmission and virulence (Figure 3, green). This is in line with the expectations of Frank (1992) and Anderson & May (1982) (2) who comment that a large virulence is favoured when considering only a single strain. The framework we adapted here can be extended for coinfection by multiple genotypes (13), however here we consider only one.

**Figure 3.**
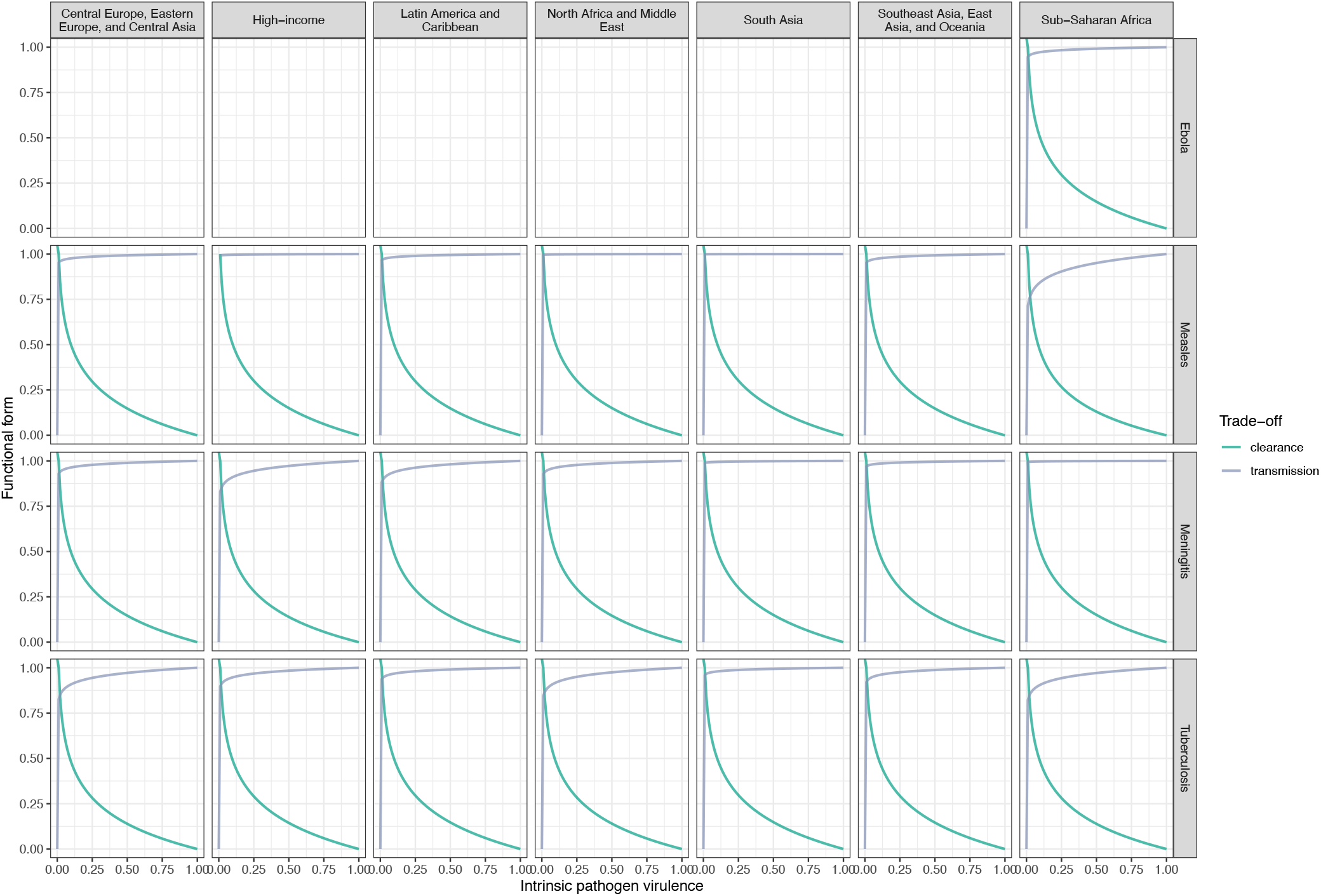
Showing the Functional form of the clearance-virulence trade off (green) and the transmission-virulence trade off (purple). These have been parameterised as 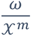 and b 𝒳^k^ respectively, where ω and b are the population-level disease specific clearance (average clearance across all ages) and disease-specific b given in Table 1. We capture the expected relationship of increasing intrinsic virulence and decreasing clearance and increasing intrinsic virulence and diminishing returns in transmission.

#### Present optimal intrinsic virulence & R_0_

Having solved for the trade-off parameter values, using Latin Hypercube Sampling we obtained the expected relationship between intrinsic pathogen virulence and *R*_0_, where an increase in virulence eventually results in diminishing returns because hosts die before they can transmit the pathogen (Figure 4). If we consider TB as an example, the age-specific susceptibility is clearly variable across regions (figure 2; bottom row). This is reflected in figure 4 (bottom row), where the optimal virulence for TB varies across regions due to this differing age structure. Optimal *R*_0_ reaches three times as large in the Central/Eastern Europe & Central Asian region as it is in the Sub-Saharan African region. Similarly Considering Meningitis, optimal *R*_0_ is twice as large in the High-Income region as it is for Sub-Saharan Africa as a function of differing age structure. The difference between present and future *R*_0_ is shown in figure S3.

**Figure 4.**
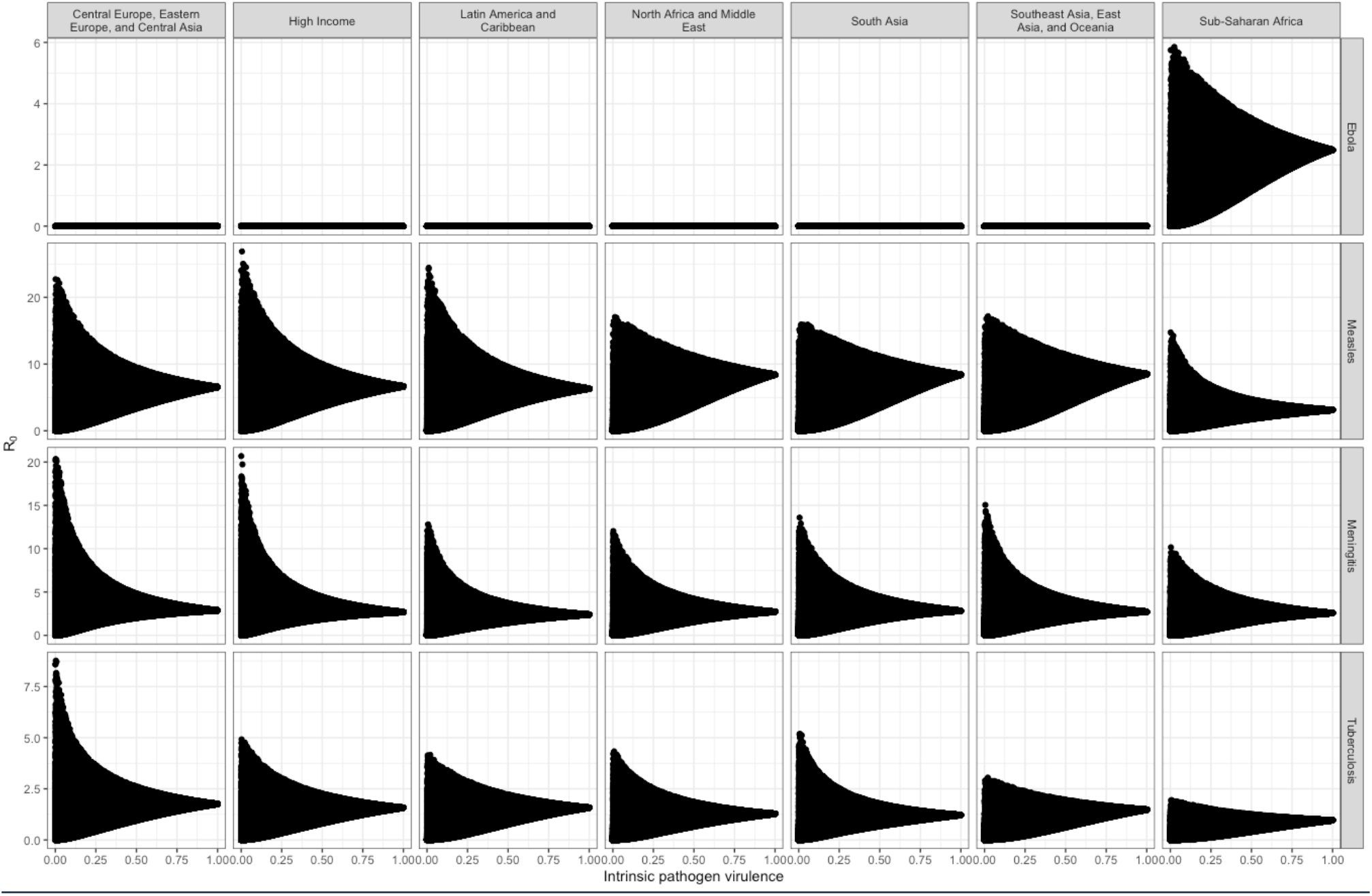
Latin hypercube sampling of the parameter space showing the relationship between intrinsic pathogen virulence 𝒳 and R_0_ parameterised with the 2017 population age structure.

#### Probability of Disease-Induced Mortality

The impact of the change in optimal virulence as a function of progressively ageing populations can be accessed via the probability of disease induced mortality *α*, here considered as the realised actual virulence *ν*𝒳 which allows for both host vulnerability to mortality and pathogen intrinsic virulence. We show that changes to population age structure occurring across all regions will impact the per case probability of mortality (Figure 5). Meningitis shows a consistent reduction in optimal virulence across all the locations, though the magnitude of this reduction is geographically dependent. Susceptibility was comparable across regions and age classes, excluding the very youngest (Figure 2), so it is likely that this variation across regions is due to changes in demographic parameters like baseline mortality because of the increasing density of old individuals with a greater force of baseline mortality (Figure S1). Alternatively, Measles noticeable variation across the regions, with an increase in optimal virulence in the Central Europe, Eastern Europe, and Central Asia region, North Africa and Middle East region, and Sub-Saharan Africa region, but a decrease in the remaining regions. It is of note that data are only available up to 60 years of for Measles, hence no predictions for older age classes. Meningitis susceptibility varies very little by region, which highlights the indirect ways in which population age-structure impacts pathogen virulence, through interactions between other population-level rates like clearance and baseline mortality rates, impacting duration of infection. Importantly, whilst a decrease in the probability of mortality can be considered a positive thing, the implication is an increase in infection duration and thus transmission duration.

**Figure 5.**
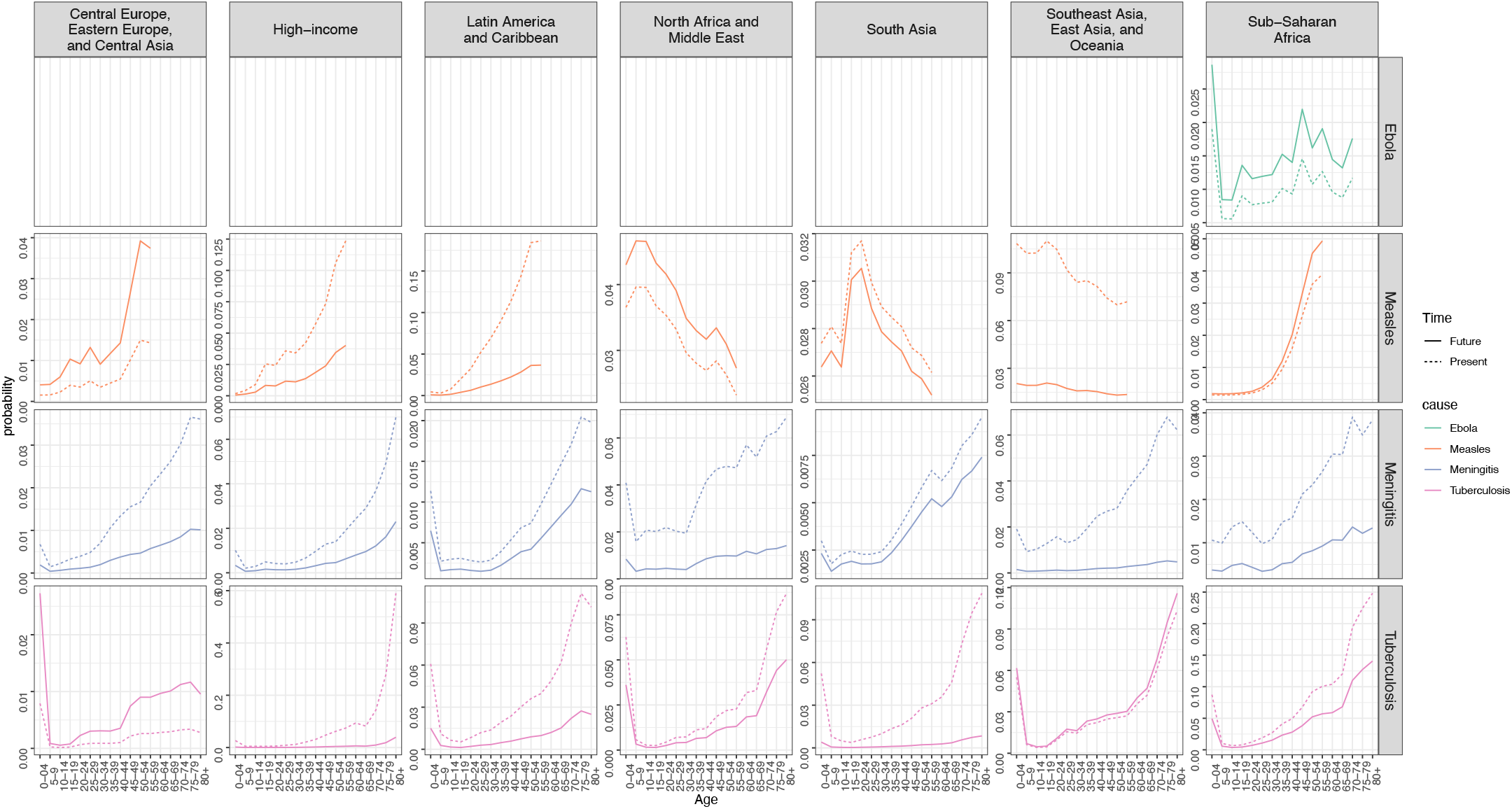
Realised actual virulence (ν𝒳) parameterised using the vulnerability ν data 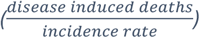 and the calculated optimal intrinsic pathogen virulence 𝒳 values for present (dotted line) and future (solid line). Pathogens are signified by colour (Ebola = green, Measles = orange, Meningitis = purple, TB = pink).

## Discussion

Here, we evidence that changing population age structure between 2017 and 2050 could drive pathogens to different virulence optima. Four communicable diseases of global interest according to the W.H.O were used to parameterise the age-specific aspects of the model. Shifts in pathogen optimal virulence were disease specific, though generally showed an upward trend. The increasing probability of mortality for many of the pathogens, as the global population ages, ultimately suggests efforts may be best spent preventing the degeneration of immune capability, rather than being left responding to it. Patterns such as these draw attention to the need for a focus on healthspan rather than lifespan from members of both biogerontological and epidemiological research. As it stands however, healthspan is largely considered in the context of non-communicable diseases, which are now commonplace as they develop over increased lifespans (21). Our results also highlight the differing needs and pressures on regional medical infrastructure and, giving an indication of how and why pressures on the health systems will develop in ageing populations.

The trade-offs between transmission and intrinsic virulence, and between clearance and intrinsic virulence, to the best of our knowledge have never been numerically estimated. There is an opportunity for this framework to be applied to more pathogens, and for these estimates to be validated experimentally. In addition, the application of this framework to pathogens that elude control despite exhaustive efforts, could provide great insight into the limitations of our interventions and new ways to limit pathogen growth based on pathogen life history optimization (22). This does, however, require high quality data. Indeed, the accurate parameterisation of this model was greatly hindered by a lack of age-specific epidemiological data.

On this note, the approximation of clearance was very general, due to a lack of quantitative data on age-specific immune capability. Ideally, clearance would have been parameterised as an age specific rate of clearance for each pathogen. This requires data on the age at infection, and continuous monitoring and recording of pathogen load for the duration of the infection. These data were not available for most pathogens, and in the rare instances when it was, the resolution was low. This lack of quantitative data on age-specific immune function is concerning, when changes in the immune system are so commonly cited as the reason for reduced resistance to infection at old age. The mechanistic parts of human immune systems have been characterised throughout life(23-25), though it would appear that these reductionist, mechanistic findings must now be brought together to quantitatively describe whole biological systems. This would also provide a unique opportunity to delineate the difference between age-specific host responses and pathogen induced host damage. In macro-parasitology for example, there is growing evidence that infection intensity does not explain the severity of infection induced morbidity(26).

The assumptions that there would be no other demographic or epidemiological changes to parameters such as baseline mortality rate, incidence, or changes in pathogen induced mortality rates occurring between 2017 and 2050 were suitable for the simplification of the model. Indeed, rates of mortality have recently plateaued in many countries, bucking the trend of globally reducing rates of mortality seen in previous years(27). However, the use of large-scale rates and population age structure to numerically parameterise the model overlooks important ecological differences in pathogen distribution, the environments in which transmission occurs, and the fine-scale nuances of R_0_. For example, the contact rates used here were chosen to produce R_0_ values that fell within the boundaries found in review articles(9-12, 28, 29), not specifically for each region. However, it is probable that the contact rate and therefore the R_0_ will vary considerably given sociodemographic determinants – for example, TB in resource limited settings with crowded living conditions (for example southern India, included in South Asia here; Ma et al 2018), will have a higher contact rate, potentially increasing R_0_ above that expected in the high-income group.

To conclude, our work suggests that shifts in population age structure will impact demographic and epidemiological parameters that could impact the evolution of pathogen virulence. The classic transmission – virulence trade-off, and recovery-transmission trade-off were numerically parameterised for the first time, highlighting the idiosyncrasies of the trade-off in the ecology of each pathogen. This work highlights the need for healthspan to be considered in the context of infectious disease, as well as the need for more detailed age-specific epidemiological data.

## Supporting information captions

Figure S1. Differences in age-specific population-level baseline mortality as a function of changing population age structure. Here we multiply age specific baseline normalised mortality estimates from 2017 by N the proportion of each age class, and the same for 2050.

Figure S2. Age, disease, and region-specific disease induced mortality.

Table S1. Mean values and standard deviations obtained from repeated simulations to calculate intrinsic virulence and R_0_ values

Figure S3. Present and Future calculated R_0_ value.

## References

1. He W, Goodkind D, Kowal P, editors. An Aging World : 2015 International Population Reports 2016.

2. Anderson RM, May RM. Coevolution of hosts and parasites. Parasitology. 1982;85:411–26.

3. Nesse RM. Evolution: medicine’s most basic science. Lancet. 2008;372:S21–S7.

4. Cousineau SV, Alizon S. Parasite evolution in response to sex-based host heterogeneity in resistance and tolerance. Journal of Evolutionary Biology. 2014;27:2753–66.

5. Cressler CE, Mc LD, Rozins C, J Vdh, Day T. The adaptive evolution of virulence: a review of theoretical predictions and empirical tests. Parasitology. 2016;143(7):915–30.

6. Boots M, Mealor M. Local Interactions Select for Lower Pathogen Infectivity. Science. 2007;315:1286-.

7. Ubeda F, Jansen VA. The evolution of sex-specific virulence in infectious diseases. Nat Commun. 2016;7:13849.

8. Rohani P, Zhong X, King AA. Contact Network Structure Explains the Changing Epidemiology of Pertussis. Science. 2010;330(6006):982–5.

9. Guerra FM, Bolotin S, Lim G, Heffernan J, Deeks SL, Li Y, et al. The basic reproduction number (R0) of measles: a systematic review. Lancet Infect Dis. 2017;17(12):e420–e8.

10. Prasinou AK. Using mathematical models to evaluate and inform immunisation strategies with MenAfriVac in the African meningitis belt. Cambridge: University of Cambridge, Darwin College; 2019.

11. Ma Y, Horsburgh CR, White LF, Jenkins HE. Quantifying TB transmission: a systematic review of reproduction number and serial interval estimates for tuberculosis. Epidemiol Infect. 2018;146(12):1478–94.

12. Van Kerkhove MD, Bento AI, Mills HL, Ferguson NM, Donnelly CA. A review of epidemiological parameters from Ebola outbreaks to inform early public health decision-making. Sci Data. 2015;2:150019.

13. Frank SA. A kin selection model for the evolution of virulence. Proceedings of the Royal Society B-Biological Sciences. 1992;250:195–7.

14. Anderson RM, Medley GF. Community control of helminth infections of man by mass and selective chemotherapy. Parasitology. 1985;90(04).

15. Gagneux S. Host-pathogen coevolution in human tuberculosis. Philosophical Transactions of the Royal Society B: Biological Sciences. 2012;367(1590):850–9.

16. Frank SA, Bush RM. Barriers to antigenic escape by pathogens: trade-off between reproductive rate and antigenic mutability. BMC Evolutionary Biology. 2007;7(1).

17. Meyers LA, Levin BR, Richardson AR, Stojiljkovic I. Epidemiology, hypermutation, within-host evolution and the virulence of Neisseria meningitidis. Proc Biol Sci. 2003;270(1525):1667–77.

18. Sofonea MT, Aldakak L, Boullosa LF, Alizon S. Can Ebola Virus evolve to be less virulent in humans? bioRxiv. 2017; Peer Community In Evolutionary Biology.

19. Carnell R. Package ‘LHS’ 2022 [

20. R Core Team. R: A language and environment for statistical computing. R Foundation for Statistical Computing. Vienna, Austria 2018.

21. Crimmins EM. Lifespan and Healthspan: Past, Present, and Promise. Gerontologist. 2015;55(6):901–11.

22. Ross A, Inobaya M, Olveda R, Chau T, Olveda D. Prevention and control of schistosomiasis: a current perspective. Research and Reports in Tropical Medicine. 2014.

23. Montecino-Rodriguez E, Berent-Maoz B, Dorshkind K. Causes, consequences, and reversal of immune system aging. J Clin Invest. 2013;123(3):958–65.

24. Hasselquist D, Nilsson J-A. Maternal transfer of antibodies in vertebrates: trans-generational effects on offspring immunity. Philosophical Transactions of the Royal Society B: Biological Sciences. 2009;364:51–60.

25. Simon AK, Hollander GA, McMichael A. Evolution of the immune system in humans from infancy to old age. Proceedings of the Royal Society B: Biological Sciences. 2015;282.

26. Wiegand RE, Secor WE, Fleming FM, French MD, King CH, Deol AK, et al. Associations between infection intensity categories and morbidity prevalence in school-age children are much stronger for Schistosoma haematobium than for S. mansoni. PLoS Negl Trop Dis. 2021;15(5):e0009444.

27. Dicker D, Nguyen G, Abate D, Abate KH, Abay SM, Abbafati C, et al. Global, regional, and national age-sex-specific mortality and life expectancy, 1950–2017: a systematic analysis for the Global Burden of Disease Study 2017. The Lancet. 2018;392(10159):1684–735.

28. Nsubuga RN, White RG, Mayanja BN, Shafer LA. Estimation of the HIV basic reproduction number in rural south west Uganda: 1991-2008. PLoS One. 2014;9(1):e83778.

29. Lo Presti A, Vacca P, Neri A, Fazio C, Ambrosio L, Rezza G, et al. Estimates of the reproductive numbers and demographic reconstruction of outbreak associated with C:P1.5-1,10-8:F3-6:ST-11(cc11) Neisseria meningitidis strains. Infect Genet Evol. 2020;84:104360.

